# Interrogation of the integrated mobile genetic elements in gut-associated *Bacteroidaceae* with a consensus prediction approach

**DOI:** 10.1101/2021.09.02.458807

**Authors:** Danielle E. Campbell, Joseph R. Leigh, Ted Kim, Whitney E. England, Rachel J. Whitaker, Patrick H. Degnan

## Abstract

Exploration of mobile genetic element (MGE) diversity and relatedness is vital to understanding microbial communities, especially the gut microbiome, where the mobilization of antibiotic resistance and pathogenicity genes has important clinical consequences. Current MGE prediction tools are biased toward elements similar to previously-identified MGEs, especially tailed phages of proteobacterial hosts. Further, there is a need for methods to examine relatedness and gene sharing among MGEs. We present VICSIN, a consensus approach for MGE prediction and clustering of predictions to provide classification. Testing of VICSIN on datasets of *Pseudomonas aeruginosa* and *Bacteroides fragilis* genomes suggests VICSIN is the optimal approach to predict integrated MGEs from poorly-explored host taxa, because of its increased sensitivity and accuracy. We applied VICSIN to a dataset of gut-associated *Bacteroidaceae* genomes, identifying 816 integrated MGEs falling into 95 clusters, most of which are novel. VICSIN’s fast and simple network-building scheme revealed a high degree of gene sharing within and between related MGE clusters. Shared gene functions across MGEs include core mobilization functions and accessory gene content, such as type VI secretion systems and antibiotic resistance genes. The MGEs identified here encode a large portion of unknown gene content, emphasizing the fact that the full diversity of MGEs and the factors they encode remain very poorly understood. Together, this work motivates more exploration of the gut mobilome, which is likely one of the most potent drivers of microbial evolution in the human microbiome.

**IMPORTANCE:** Mobile genetic elements (MGEs), including phages and integrative and conjugative elements (ICEs), drive the diversity and function of microbial communities through horizontal gene transfer. Current tools to predict MGEs in genomic sequence data are highly focused on phages, and are biased against the discovery of novel MGEs. We present VICSIN, a consensus approach to MGE prediction that is able to find a diversity of MGEs, particularly in poorly-understood bacterial taxa. By applying VICSIN to a large database of diverse *Bacteroidaceae* genomes, we have been able to get a distinct view of the gut mobilome, extending beyond the phageome. These novel MGEs belong to related groups, sharing a significant amount of functional gene content within and between groups, supporting a mosaic model of evolution for ICEs. Understanding how phages evolve in *Bacteroidaceae* hosts, however, remains elusive and highlights the need for more experimental research.

## INTRODUCTION

Horizontal gene transfer (HGT), the sharing of DNA between co-existing organisms, is a primary mechanism shaping microbial communities, and a driver of genomic plasticity and adaptation (1–5). Across microbial communities, it is postulated that the vast majority of HGT is mediated by mobile genetic elements (MGEs) (4–7). MGEs encode their own mechanisms that allow for their transfer between and their maintenance within cellular hosts, thereby circumventing two of the main barriers to HGT: cell entry and persistence (4). HGT occurs at especially high rates in the human gut microbiome (8, 9), which has been attributed to both the high density of microbes (10, 11) and the activity of some MGEs to host-associated conditions, such as inflammation (12, 13). MGE activity has important consequences for all microbial hosts and communities, including in the human gut. Within the gut microbiome, MGEs disseminate antibiotic resistance genes, virulence factors, and other advantageous traits, both in symbiotic and pathogenic contexts (3, 14–17). Characterizing MGE diversity is crucial for understanding how genetic diversity, genes of interest, and relevant traits are maintained within and disseminated between microbes.

The process of characterizing MGE diversity presents two major challenges: (i) accurate and sensitive detection of MGEs, and (ii) MGE classification. An increasingly large number of tools have been developed to search genomic sequence data for putative MGEs using distinct computational approaches (18). Each of these individual approaches to MGE prediction has advantages and limitations. Similarity-based search tools are biased towards the identification of MGEs similar to previously characterized MGEs, and therefore perform poorly at identifying novel MGEs (18, 19). *De novo* prediction tools use generic models of MGE sequence characteristics but exhibit variable accuracy and sensitivity depending on the microbial host (18, 19). Genome alignment tools identify all non-core genomic elements, which include MGEs but can also include other accessory genomic features, such as recombination hotspots (20), cell surface modification loci (20, 21), and CRISPR-Cas systems (22). Machine learning and deep learning-based tools may require large dataset inputs or customized training data, which may not be available for most microbial systems (18, 23, 24). Furthermore, MGE identification is most thoroughly tested on clinically-relevant and well-studied host microbes, especially those in the phylum *Proteobacteria* (19, 23–27), and is largely tailored to identifying integrated viruses (*i*.*e*., prophages), neglecting the numerous other types of MGEs (18, 28). Although testing these tools on lesser-understood microbial hosts and MGEs is intrinsically difficult, the current lack of representation across host and MGE diversity has resulted in biases in MGE prediction (27).

The second major challenge to understanding MGE diversity is understanding how they are related, and their further classification into related groups. Although many types of MGEs have been identified, we will focus on viruses and integrative and conjugative elements (ICEs), which appear to be the predominant means of HGT in the gut microbiome (29). Traditionally, viral classification has been accomplished by the International Committee for the Taxonomy of Viruses (ICTV), which relies on a combination of viral characteristics to classify new viruses, including viral particle morphology, genome composition, and homology (30). There is no universally accepted method for the classification of most other MGEs. One method for ICE classification is based on integrase gene homology and synteny of ICE backbone features (31), however the mosaic nature of ICEs suggests classification based heavily on a single gene may not be accurate (32). The mosaicism within ICEs and across all MGE types is a result of recombination events between otherwise unrelated MGEs coexisting within a single host cell (32, 33). While the continuous nature of MGE diversity is perhaps the biggest challenge to their classification, it also calls for a generalizable classification scheme that does not currently exist, in order to better understand their diversity and relatedness.

The genus *Bacteroides* represents one of the most common and abundant microbes associated with the human gut (34), having coevolved with hominid hosts for millions of years (35). *Bacteroides* genomes encode diverse suites of accessory genomic elements including polysaccharide utilization loci (36, 37), capsular polysaccharide synthesis loci (38), CRISPR-Cas systems (39, 40), and a diversity of integrated MGEs (41, 42). Although *Bacteroides* species engage in beneficial interactions with hosts (*e*.*g*., production of short chain fatty acids, immune system development), they can also be opportunistic pathogens outside and inside the gut lumen. *Bacteroides* species commonly carry ICEs encoding antibiotic resistance genes (ARGs) that can be transferred among symbiotic and pathogenic microbes in the gut, thus acting as important antibiotic resistance reservoirs (15, 43, 44). Further, enterotoxigenic *B. fragilis* strains secrete hemolysins and toxins associated with inflammation and colon cancer, some of which are carried by ICEs (45–48). Given their clinical relevance, genomic diversity, and large amount of sequence data, the *Bacteroides* presents an ideal case study for examining MGE diversity.

Here we describe a new tool: Viral, Integrative, & Conjugative Sequence Identification & Networking (VICSIN). VICSIN uses a consensus approach to combine similarity-based searching, *de novo* prediction, and genome alignment for integrated MGE prediction. VICSIN then classifies MGEs using nucleotide homology to network and cluster them into related groups. We test VICSIN using two distinct benchmark datasets: *Pseudomonas aeruginosa*, a very well-understood microbial host of diverse MGEs from the phylum *Proteobacteria*, and *Bacteroides fragilis*, a lesser-studied organism from the divergent phylum *Bacteroidetes*. Benchmark analyses suggest that although VICSIN performs comparably to its constituent tools for *P. aeruginosa* genomes, VICSIN is more accurate and sensitive for the *B. fragilis* benchmark dataset. Thus, we suggest VICSIN is a superior tool for MGE discovery in poorly characterized hosts. We applied VICSIN to a large dataset of *Bacteroides* genomes, revealing a highly diverse set of related MGEs that has gone unexplored.

## RESULTS

### VICSIN predicts and networks integrated MGEs

We developed VICSIN as a tool for (i) comprehensive prediction of MGEs integrated in microbial genomes and (ii) clustering of MGEs as a method of classification (Fig. 1). VICSIN uses a consensus approach to MGE prediction, utilizing the pre-existing MGE prediction tools VirSorter (19), PhiSpy (25), and AGEnt (26). Further, VICSIN integrates optional nucleotide BLAST steps with user-input databases of CRISPR spacers and/or known MGEs. Consensus boundaries of overlapping predictions are determined for each genome. These consensus regions are then compared across the set of analyzed genomes using nucleotide BLAST and boundaries are extended if they are sufficiently similar to an independent consensus prediction (“Re-BLAST”), producing the final VICSIN predictions. Finally, VICSIN predictions are networked on the basis of percent length aligned (PLA) using reciprocal nucleotide BLAST. MGE pairs with PLAs greater than 20% are passed to MCL for clustering (Fig. 1).

**Figure 1.**
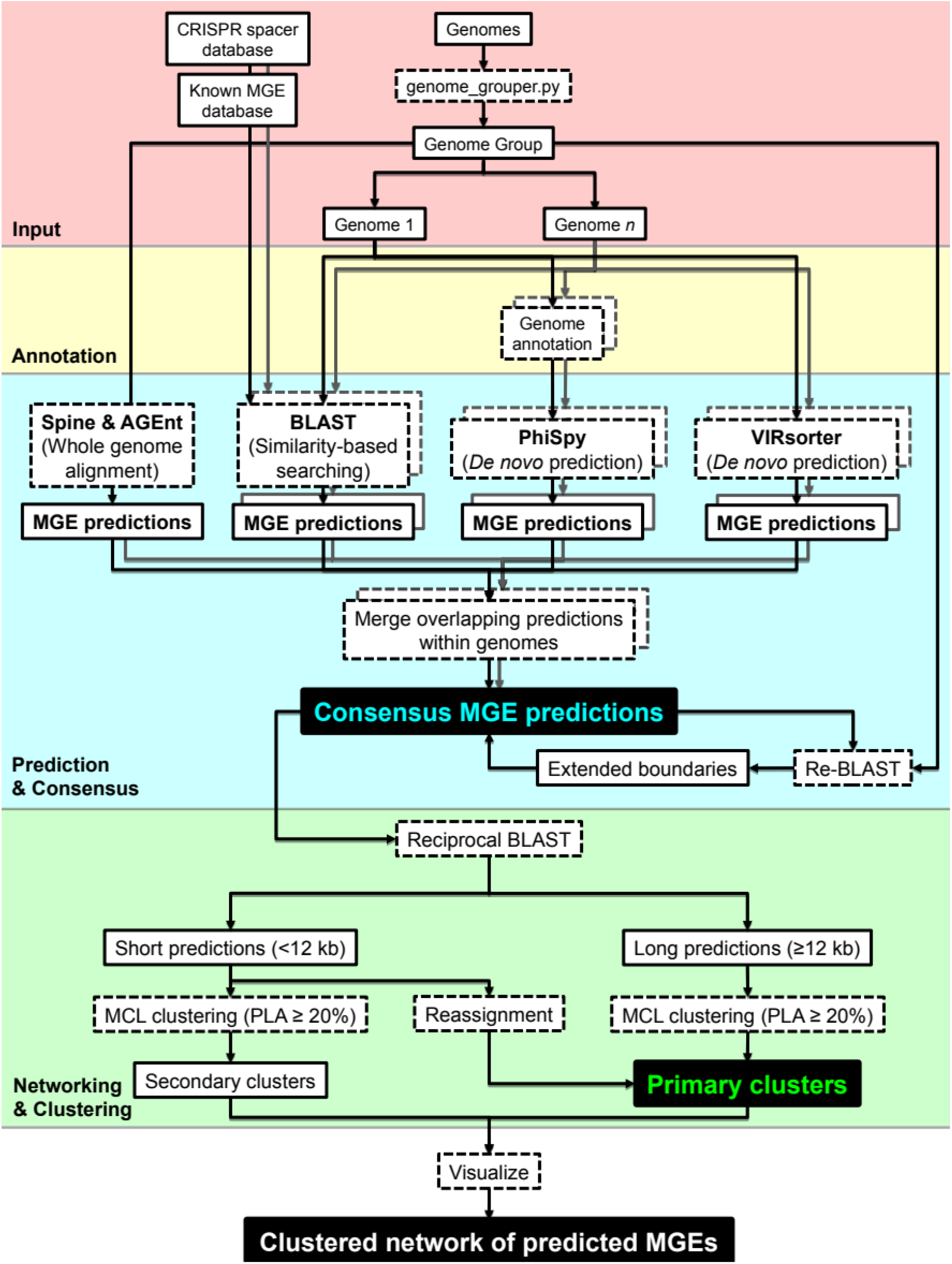
Overview of the VICSIN process. VICSIN takes groups of closely related microbial genomes as the only required input; CRISPR spacer and known MGE databases are optional inputs. VICSIN runs each of its constituent prediction tools: Spine and AGEnt, PhiSpy, VirSorter, and nucleotide BLAST (optional). Consensus predictions are determined and their boundaries may be extended in the Re-BLAST step. MGE predictions are split into two groups by their length. Reciprocal nucleotide BLAST is used to calculate percent length aligned (PLA), which is the basis for MCL clustering. Network and cluster outputs are formatted for viewing with external network visualization software. Dashed outline boxes indicate processes; solid outline boxes indicate input or intermediate result data; black filled boxes indicate primary result outputs.

VICSIN is designed to take groups of ≥5 related bacterial genomes as input. To test VICSIN, two benchmark datasets were developed for the divergent bacterial species *P. aeruginosa* and *B. fragilis* (Fig. 2). Each benchmark dataset is comprised of 6 closely related genomes (Fig. 2). In order to prevent drops in prediction precision, VICSIN requires input genomes be grouped below the species level, using a cutoff of ≥70% PLA by Spine, approximately corresponding to >98% average nucleotide identity (ANI) in both benchmark datasets (Fig. S1). For both datasets, “known MGEs” integrated in each genome were identified based on literature reports or manual annotation (Table S1), identifying 40 known MGEs in the *P. aeruginosa* genomes, and 38 known MGEs in the *B. fragilis* genomes. Though VICSIN can use known MGEs for prediction by BLAST, this option was not used for benchmarking purposes.

**Figure 2.**
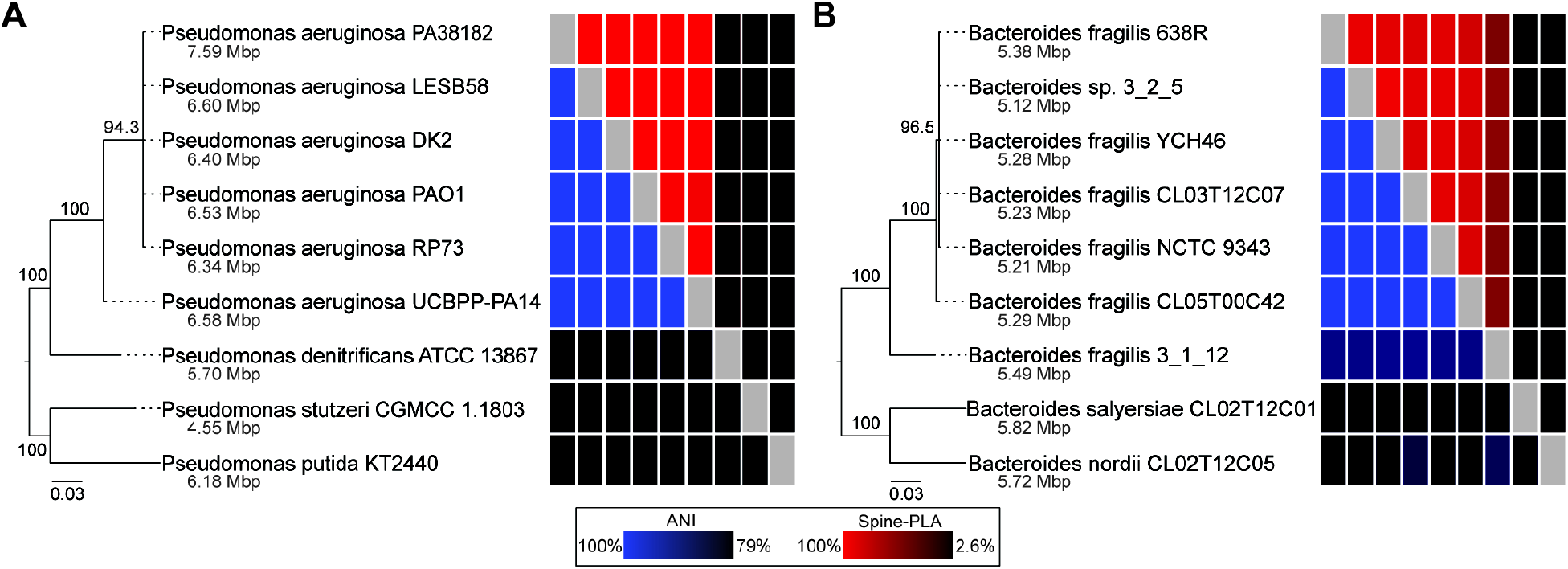
Benchmark datasets for testing VICSIN. Benchmark datasets were constructed from genomes of (A) *P. aeruginosa* and (B) *B. fragilis*. Phylogenies for each dataset were estimated by maximum likelihood from concatenated alignment of 13 core genes. Support values above tree branches represent bootstraps from 100 replications. Adjacent heatmaps depict average nucleotide identity (ANI) (49) and PLA calculated by Spine (Spine-PLA) for each genome pair except for self-pairs. Each dataset includes one intermediate genome (*P. denitrificans* ATCC 13867 and *B. fragilis* 3_1_12) and two outgroup genomes (*P. stutzeri* GCMC 1.1803, *P. putida* KT2440, *B. salyersiae* CL02T12C01, and *B. nordii* CL02T12C05), which were used to test the effects of genome relatedness on VICSIN accuracy and sensitivity (Fig. S1). Genome sizes are indicated for each genome in grey.

Using the known MGEs in these two benchmark datasets, we compared the accuracy and sensitivity of VICSIN to its individual constituent tools (Fig. 3). We used four main metrics to evaluate the effectiveness of the tools: precision (*i*.*e*., the number of true positive predictions, divided by the total number of predictions), recall (*i*.*e*., the number of true positive predictions, divided by the total number of known MGEs), coverage (*i*.*e*., the percent length of a known MGE covered by a true positive prediction), overprediction (*i*.*e*., the excess length of a true positive prediction that does not cover a known MGE), and total false positive length (*i*.*e*., the sum of the lengths of all false positive predictions in a genome). True positive predictions were defined to be any prediction overlapping a known MGE in a benchmark genome.

**Figure 3.**
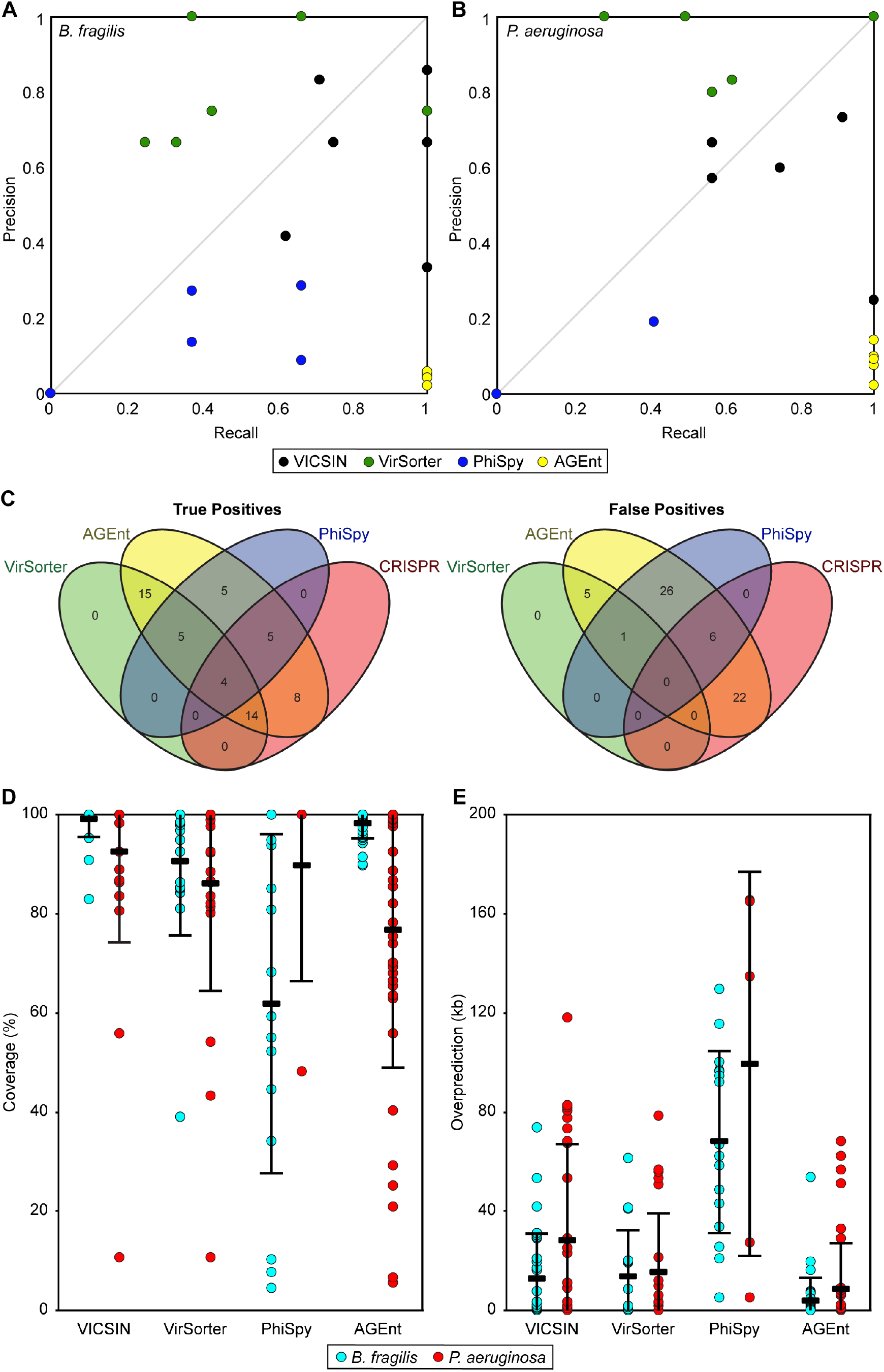
MGE prediction with VICSIN is more sensitive and accurate than with its constituent tools. MGE prediction precision and recall were calculated for each of the six genomes from the (A) *B. fragilis* and (B) *P. aeruginosa* benchmark datasets. Precision was calculated as the proportion of true positive predictions to total predictions made. Recall was calculated as the proportion of true positive predictions to known MGEs. (C) The constituent tools (VirSorter, AGEnt, PhiSpy, CRISPR BLAST) comprising true positive (n = 56) and false positive (n = 60) MGE predictions were summed for both *B. fragilis* and *P. aeruginosa* datasets. Except for VirSorter predictions, VICSIN requires consensus between two methods to accept a prediction. (D) For each in-group known MGE with a corresponding prediction, percent coverage was calculated as the ratio of the correctly predicted basepairs to the actual total basepairs of the known MGE. The black bars represent averages and standard deviations are displayed by error bars. (E) For each in-group known MGE with a corresponding prediction, the length of overprediction was calculated and each represented as a colored circle. Averages are displayed as black rectangles and standard deviations are displayed by error bars.

Analysis of the *B. fragilis* dataset found that VICSIN is more accurate and sensitive than its constituent tools VirSorter, PhiSpy, and AGEnt for all six genomes (Fig. 3A). For some of the *P. aeruginosa* genomes (*i*.*e*. strains DK2, PAO1, and RP73), VirSorter has greater precision and recall than VICSIN (Fig 3B). Inspection of these predictions revealed that the dropoff in VICSIN’s accuracy was primarily driven by PhiSpy and AGEnt (Fig. 3C), an expected result given the low precision of PhiSpy (Fig. 3A, 3B), and that AGEnt detects all variable components of microbial genomes (26). However, 28 of 60 (47%) false positive predictions were supported by CRISPR BLAST matches, suggesting there may remain a number of unrecognized MGEs in the benchmark genomes. Overall, VICSIN predictions exhibit high coverage of known MGEs (Fig. 3D) and low overprediction (Fig. 3E), especially in the *B. fragilis* dataset. Together, this analysis suggests VICSIN is at least as accurate and sensitive as its constituent tools for *P. aeruginosa* host genomes, and exceeds the accuracy and precision of those constituent tools for *B. fragilis*. This may be a result of *P. aeruginosa* genomes being key elements in the testing and training of VirSorter, PhiSpy, and AGEnt [19, 25, 26]. We suggest VICSIN is the optimal approach to MGE prediction in poorly-characterized host genomes, especially those outside the phylum *Proteobacteria*.

Classification of predicted MGEs is a critical step for subsequent analyses of evolutionary history, mode of transfer, and host range. The existing tools ClustAGE (50) and vConTACT 2.0 (51) use different methods to classify MGEs, each with advantages and limitations. ClustAGE divides input genomic elements into subelements (*e*.*g*. gene blocks) and bins these elements using nucleotide similarity. These subelement bins allow ClustAGE to examine the mosaic sharing of nucleotide blocks, offering a fine-scale view of relatedness, although it does not attempt cluster the MGEs as a whole. vConTACT, which was designed for viral sequence data, first clusters the predicted proteins encoded by input genetic elements, and subsequently uses these shared protein profiles to generate viral clusters. Like ClustAGE, VICSIN’s clustering does not require open reading frame annotation, translation, or protein clustering. However, instead of identifying shared subelements, VICSIN generates MGE clusters based on the extent of shared nucleotide sequence among the input MGEs. To evaluate the effect these methods have, the known MGEs in the *P. aeruginosa* benchmark dataset (Table S1) were analyzed in parallel with VICSIN, ClustAGE (50), and vConTACT 2.0 (51). From the 40 known MGEs used as input, both VICSIN and vConTACT produced the fewest clusters (Table 1), and nearly identical networks (Fig. S2A, S2B). This is expected, since both methods were designed for the similar purposes of classifying MGEs or viruses, respectively, and employ large lengths of sequence similarity. The fundamentally different approach of ClustAGE results in a large number of subelement bins, illustrating the high degree of connectivity among the known MGEs over short sequence lengths; however, it does little to provide a means of classifying the MGEs more broadly (Fig. S2C).

**Table 1.** Clustering of *P. aeruginosa* benchmark MGEs.

|  | VICSIN | vConTACT 2.0 | ClustAGE<sup>a</sup> |
| --- | --- | --- | --- |
| No. of clusters/bins | 6 | 6 | 83 |
| Largest cluster/bin | 6 | 6 | 6 |
| No. singletons | 23 | 24 | 67 |

<sup>a</sup>ClustAGE does not generate clusters; bins were counted instead.

Importantly, VICSIN offers several practical advantages while achieving similar results to vConTACT. Prior to running vConTACT the user must also predict protein encoded on their MGEs, and then generate protein clusters using programs in the vConTACT toolbox. This process, however, is not designed to handle multiple MGEs and requires additional work to reformat and compile protein data inputs. VICSIN eliminates these additional steps by operating directly on the nucleotide sequences. Further, by using nucleotide data, VICSIN avoids the possibility of protein prediction inaccuracies, as MGEs can be challenging to properly annotate (52). vConTACT and VICSIN are both built on BLAST and MCL, making them equally scalable. Because of its practical advantages and its reduced susceptibility to error and compute time during protein calling and data format conversion, we conclude that VICSIN is a superior tool for MGE clustering.

### *Application of VICSIN to characterize the* Bacteroidaceae *integrated mobilome*

A subset of genomes was selected from a database of 341 *Bacteroides* genomes accessible on NCBI or sequenced in-house (Table S2). As was performed for the benchmark datasets, genomes were compared by pairwise Spine-PLA (Fig. S3). From the similarity matrix, two groups of interest were chosen for analysis with VICSIN: (1) 61 *B. fragilis* genomes, (2) and 27 *P. vulgatus* or *P. dorei* genomes. *B. fragilis* is commonly associated with extraintestinal infections and perhaps the best-studied *Bacteroides* species (53). As a result, many known MGEs have been characterized for it, including ICEs (54–60) and phages (61–65). *P. vulgatus* and *P. dorei* constitute a clade divergent from that of *B. fragilis*, and together have fewer known MGEs (66, 67). This scheme was chosen in order to characterize MGEs both within host groups and also to sample across the diverse *Bacteroidaceae* family. The *B. fragilis* and *P. vulgatus* groups were arbitrarily split into subgroups of 7-10 genomes, in order to keep AGEnt sensitivity high and compute times low. Because VICSIN prediction does not scale, each subgroup was run independently for prediction, and those predictions were later aggregated for VICSIN MGE cluster analysis.

VICSIN identified a total of 816 MGEs across the 88 host genomes (Table S3). This constitutes 42.3 Mb of predicted mobile sequence out of 468 Mb of total genome length (9.0%). This is greater than the mobile content of the initial *P. aeruginosa* (4.3%) and *B. fragilis* (5.0%) benchmark datasets. This may be the result of false positive predictions, reflecting other accessory genomic elements. Therefore, we recommend manual examination of predictions, as for all MGE prediction tools, especially from large datasets (≥50 genomes). However, it should be noted that 9% of a genome consisting of MGEs is well within the range what has been observed for other organisms (68, 69). Across the 816 VICSIN predictions, 96% were supported by AGEnt, 50% by VirSorter, 38% by CRISPR match, and 31% by PhiSpy (Fig. S4).

All 816 MGE predictions were pooled with a set of known *Bacteroidaceae* MGEs including phages and ICEs (Table S4). These predicted and known *Bacteroidaceae* MGEs were clustered with VICSIN, generating 95 MGE clusters that represented 770 individual MGE predictions and 41.1 Mb of predicted mobile sequence (∼97% of predicted MGEs) (Fig. 4). Only 46 predicted MGEs were determined to be singletons, having ≤20% PLA with the other MGEs. Each MGE was assigned an MGE type (*i*.*e*. phage, ICE, mobile, or unknown) on the basis of Pfam annotations (Table S3, S5).

**Figure 4.**
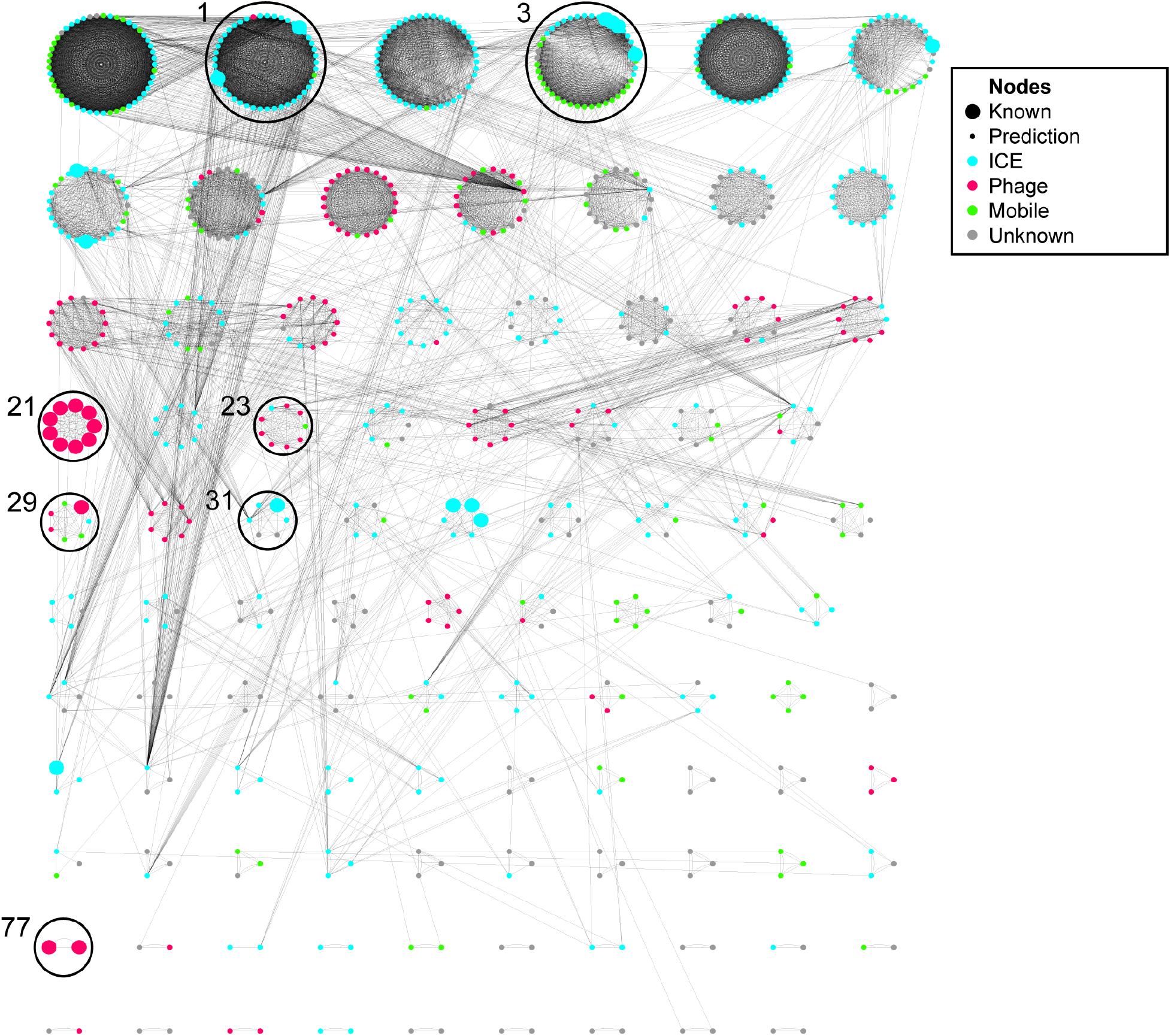
Network of clustered MGEs predicted from *Bacteroidaceae* hosts. VICSIN predicted MGEs from *B. fragilis* and *P. vulgatus* were clustered using VICSIN with a set of known *Bacteroidaceae* MGEs (Table S4). Each node nodes represents an MGE. Node size indicates if an MGE is a prediction (small) or known (large); node colors correspond to the MGE type, either based on known type or on Pfam assignment. Edges are drawn between nodes when PLA ≥ 20%. Clusters 1, 3, 21, 23, 29, 31, and 77 are indicated.

### Genetic recombination in VICSIN-predicted phage clusters

It was observed that three clusters with known phages are isolated in the network having low/no PLA above 20% with other MGEs (Fig. 4, S4). This includes the crAssphages (Cluster 21), Hankyphage and its putative relatives (Cluster 29), and two siphophages (Cluster 77), but not *Salyersviridae* temperate phages (Cluster 23). Given the low connectivity of some phage clusters in the network analysis, it suggests that the rate of recombination between MGEs may vary by MGE type or cluster. Notably, this pattern is not determined by phage lifestyle (*i*.*e*. obligately lytic versus temperate). Hankyphage is a Mu-like phage capable of being stably integrated, but we did not identify sufficient shared DNA sequences with other MGEs. In contrast, a recently-described group of temperate phages from the proposed family *Salyersviridae* does exhibit some gene sharing with other MGE clusters. Further research will be needed to determine the mechanisms affecting recombination rates between specific MGE types. While further sampling of MGEs may find more evidence of gene sharing between phages and other MGEs in *Bacteroidaceae* hosts, these results suggest it is less common, and certainly much rarer than the rate of gene sharing among ICE-like MGEs (Fig. 4).

### Encoded protein functions of VICSIN-predicted MGEs

We hypothesized that the gene content uniting an MGE cluster would be comprised of core genes encoding the essential functions for MGE transmission and persistence, and the gene content shared between MGE clusters would be comprised of accessory genes that benefit the *Bacteroidaceae* host. To examine gene sharing at a functional level, predicted MGE genes were annotated using Pfams (Fig. 5).

**Figure 5.**
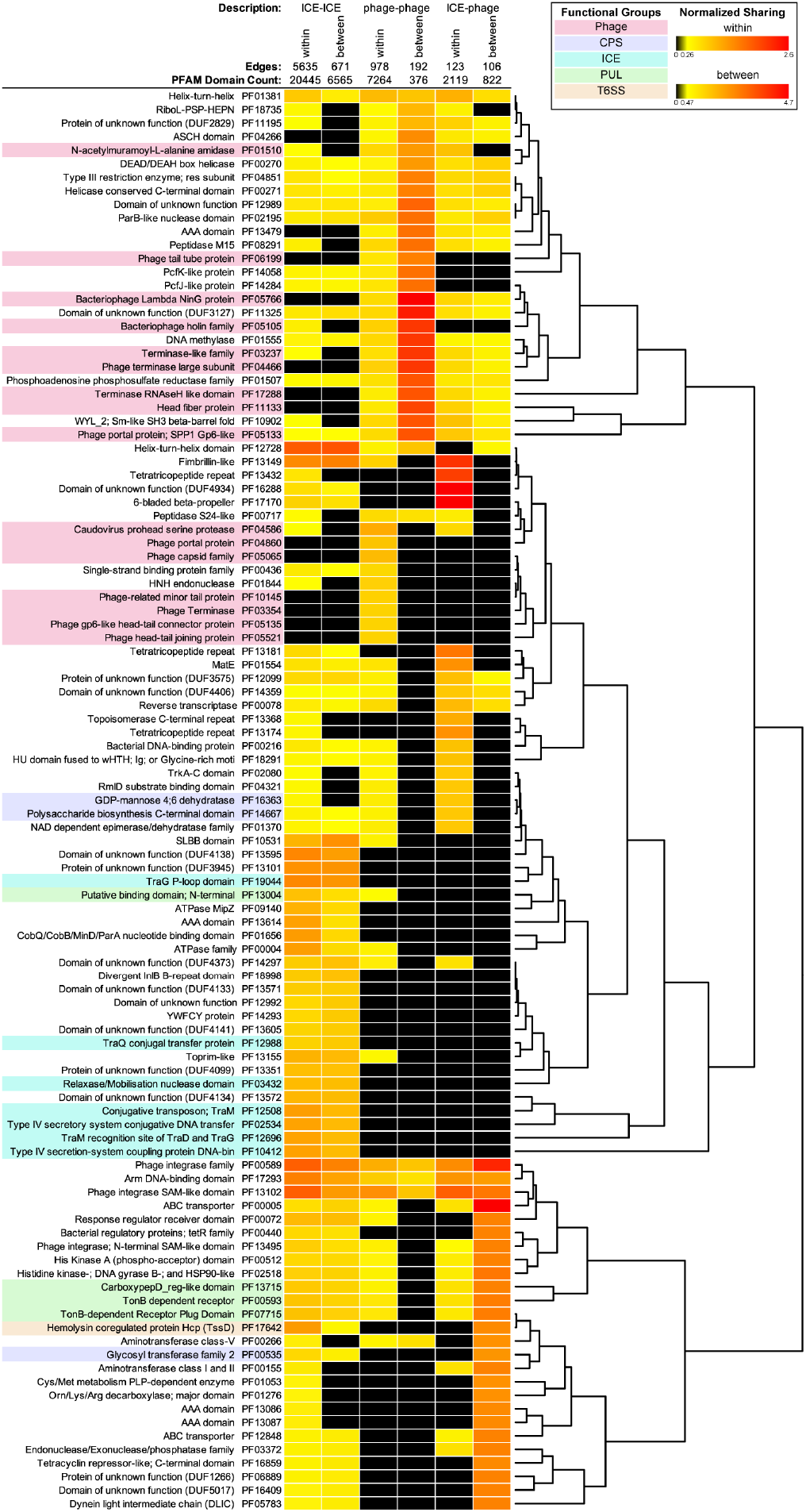
Analysis of functional gene sharing within and between MGE clusters using Pfams. Genes which were shared between MGE predictions were annotated using the Pfam database and assigned to a category on the basis of the MGE types involved, and whether that gene is shared within or between clusters. Mobile and unknown MGE categories are not shown. The number of sharing events for each Pfam was normalized to the total number of Pfams counted in each category (Pfam Domain Count). 105 Pfams are shown, representing the 25 most frequently shared Pfams in each category. Pfams were assigned to functional groups based on the genetic elements they are known to be associated with; ambiguous Pfams were not assigned to a group. Functional groups highlighted: phage-associated (pink), capsule synthesis (blue), ICE-associated (cyan), carbohydrate utilization (green), and T6SS (orange). Rows were ordered and the dendrogram were generated using heatmaps.2 in the gplots package in R (https://www.rdocumentation.org/packages/gplots/versions/3.0.4/topics/heatmap.2).

This analysis supports our hypothesis that sharing within MGE clusters would be defined by core transmission genes. Many of the most frequently shared genes between ICEs within a cluster (ICE-ICE within) are those that encode the Type IV secretion system (T4SS). Likewise, some the most frequently shared between phages within a cluster (phage-phage within) encode structural components of the phage capsid, or the machinery for packaging the phage genome. Gene sharing between ICEs and phages within a cluster (ICE-phage within) is largely driven by functions not readily assignable to an MGE type, or to accessory genes like those associated with capsular polysaccharide synthesis (CPS) loci or polysaccharide utilization loci (PUL). This is reflective of the ambiguous nature of those MGE clusters apparently containing ICEs and phages, which further investigation may find are comprised of novel MGE types or chimeric MGEs.

The Pfam analysis, however, only partially supports our hypothesis that gene sharing between clusters would be driven by accessory genes. Functional gene sharing between MGE clusters is dominated by the same core genes which drive gene sharing within those clusters. This sharing of core genes between MGE clusters does support the hypothesis that recombination is a powerful contributor to MGE evolution, leading to their modular and mosaic nature. While some accessory genes, such as Type VI secretion system (T6SS) genes, are shared between clusters, their frequencies are dwarfed by that of core genes. This is likely due to several factors, including difficulty in annotating potential accessory genes using Pfams and the potential for a high diversity of accessory genes compared to core functional genes.

### Extensive synteny within VICSIN MGE clusters

The sharing of genetic content was also examined through representative clusters. Clusters of interest were chosen on the basis of (i) having a known MGE member (Fig. S5), and (ii) having members from more than one species (Fig. S6). We postulated that clusters with MGEs from more than one host species are least likely to contain false positive predictions, by virtue of being recently mobile. Two clusters each representing ICEs and phages were selected; no attempt was made to examine clusters composed of unknown MGE types, which are difficult to annotate. Cluster 1 (CTn86-like elements) and Cluster 3 (CTnDOT-like elements) were chosen as representative of ICEs (Fig. 6A, 6B), and both show extensive sharing of core functional genes. Shared genes within these clusters include the *tra* operon, which encodes the T4SS, and can include the *mob* operon, encoding the relaxosome. Further, accessory genes in ICEs (*i*.*e*. bile salt hydrolase, xylanase, antibiotic resistances) are variable within their respective clusters. Representative phage clusters were Cluster 23 (*Salyersviridae*-like elements) and Cluster 29 (Hankyphage-like elements) (Fig. 6C, 6D). Both phage clusters exhibit a greater degree of similarity within the cluster than do the representative ICE-like clusters, covering the majority of these phage genomes. This is likely because more of a phage genome can be considered core; phages encode a large number of genes involved in capsid structure and assembly, genome packaging, host lysis, DNA replication, and regulation.

**Figure 6.**
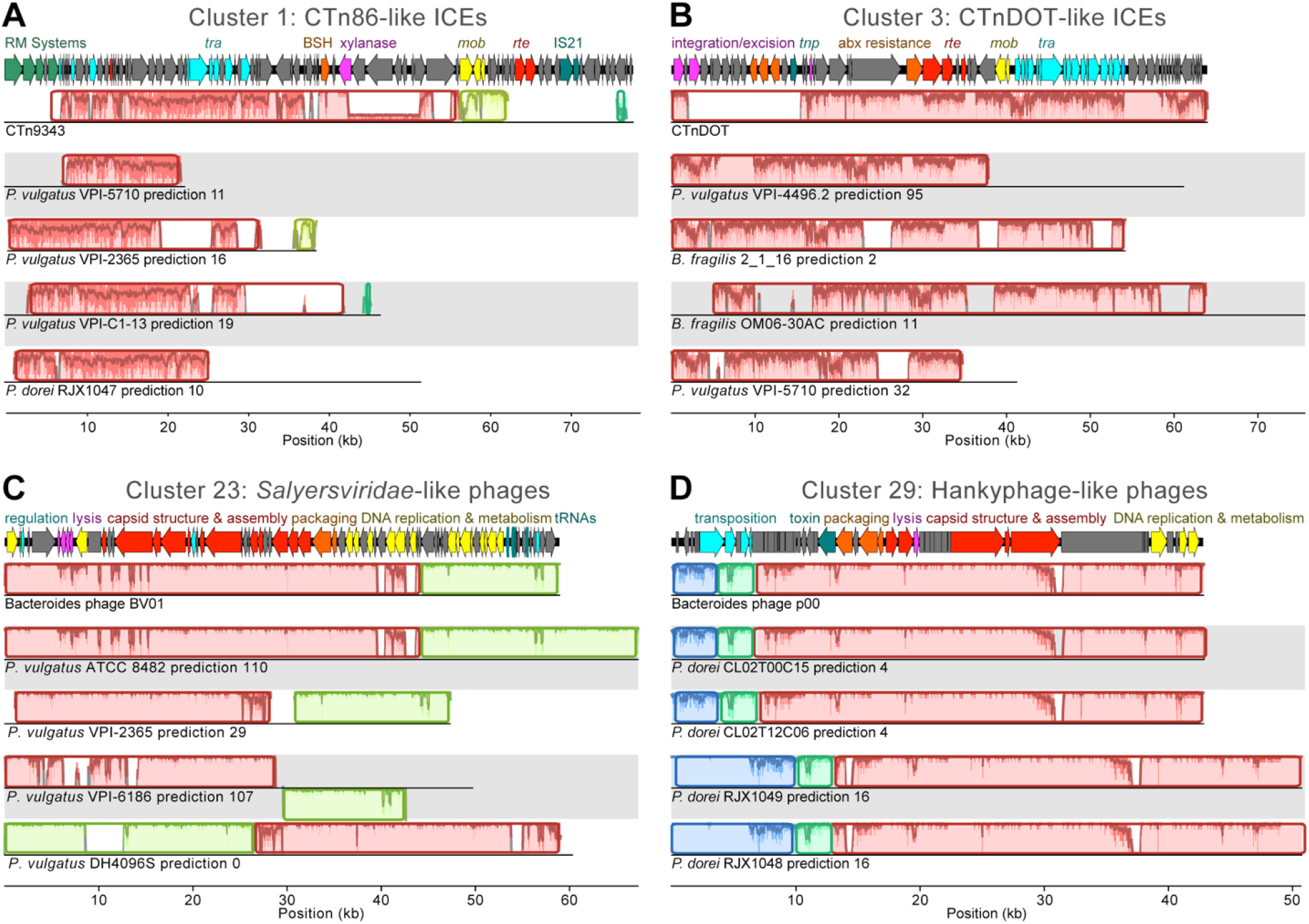
Alignment of three example VICSIN MGE clusters. Nucleotide alignments were made for select nodes from (A) Cluster 1, (B) Cluster 3, and (C) Cluster 23, and (D) Cluster 29. Alignments each show four VICSIN predictions and one known MGE: CTn9343, CTnDOT, Bacteroides phage BV01, and Bacteroides phage p00. MGE genome maps at top represent the known MGE. Alignments made in Mauve (70). Colors represent locally collinear blocks and curves represent sequence conservation.

### Detection of clinically-relevant traits in VICSIN-predicted MGEs

Finally, functional gene sharing was further interrogated by searching for specific clinically relevant functions among the VICSIN predictions. First, we examined antibiotic resistance genes (ARGs) (Fig. 7A). Twenty of the 816 VICSIN predicted MGEs were determined to encode a total 24 resistance genes. All were characterized resistance genes common among the Bacteroidetes: *cfxA3, ermF, ermG, tetX, tetQ* and *linA*, each conferring cefoxitin, clindamycin, erythromycin, tetracycline, tetracycline, and lincomycin resistance, respectively.

**Figure 7.**
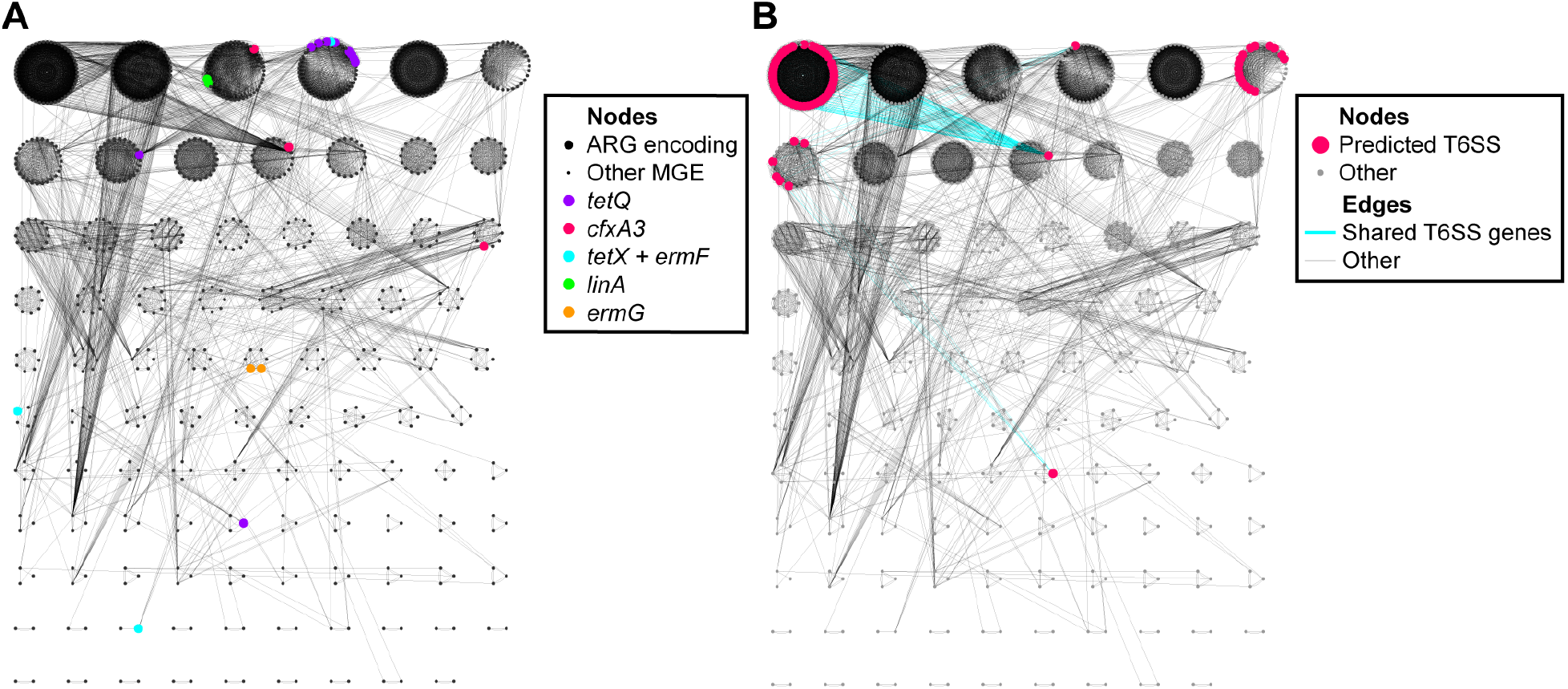
Sharing of ARGs and T6SS genes among *Bacteroidaceae* integrated MGEs. (A) ARGs from the CARD database were identified in VICSIN-predicted MGEs. (B) T6SS predictions were made for MGEs encoding *hcp, tssB*, and *vgrG*, which are conserved in all T6SS genetic architectures known to occur in *Bacteroidaceae* genomes (Table S6).

We hypothesized that antibiotic resistance genes would be associated with particular MGE clusters due to strong selection for MGEs to maintain their ARGs over evolutionary time. Yet, given the large size of the MGEs predicted (average length = 51.8 Kb), the short length of the ARGs observed (1,974 bp for the longest ARG, *tetQ*), and the 20% PLA cutoff for clustering, we did not expect the sharing of ARGs to drive MGE clustering. However, our initial hypothesis was true only for *linA*, which was detected exclusively in two MGEs in Cluster 2 (Table 2). Further sampling may reveal that *linA* is not restricted to a single MGE cluster. We determined that all of the other ARGs detected were distributed in multiple MGE clusters. The tetracycline resistance gene *tetQ* is common and highly conserved in Cluster 3, and although it is also present in two other clusters, is found in 8 out of 39 predictions (21%) within the group. This suggests that although *tetQ* is maintained over evolutionary time by some members of a cluster, stochastic loss events may cause its absence in other MGEs of the cluster. Two MGE predictions encode two copies of the same ARG (*i*.*e. tetX* in *P. vulgatus* VPI-4496.2 prediction 23 and *ermG* in *B. fragilis* OF05-13AC prediction 17), although these ARGs are not identical to one another. These may be unique alleles of the same ARG and confer resistance to the same antibiotics, or they might confer unique resistance phenotypes for which a resistance gene is not yet known. Three MGEs encode both *tetX* and *ermF* (*i*.*e. P. vulgatus* VPI-4496.2 prediction 23, *B. fragilis* CL07T12C05 prediction 72, *P. vulgatus* VPI-BV8526 prediction 108), although these shared ARGs do not correlate with their associated clusters. In two of these three MGE predictions, *ermF* and *tetX* occur in a conserved gene cassette. Together, the distribution of *tetX* and *ermF* supports the previous theory that ARGs are readily transferred individually and in cassettes between co-infecting MGEs via recombination (15, 43).

*Bacteroidaceae* MGEs are also known to carry T6SSs, which confer a competitive advantage to their hosts in the gut by antagonizing neighboring bacteria. To search for T6SSs in VICSIN predictions, three genes were used as queries: *hcp, tssB*, and *vgrG*. These genes are universal among the three T6SS genetic architectures known to occur in *Bacteroidaceae* genomes. Of the 816 VICSIN predictions, 71 were found to encode predicted T6SS. T6SSs primarily occurred in three clusters: Cluster 0, Cluster 5 (CTnBST-like ICEs), and Cluster 6 (*P. vulgatus*-associated ICEs). Evidence of sharing of these T6SS genes is apparent between clusters (*i*.*e*. between Clusters 0 and 9, and between Clusters 6 and 52), suggesting they may also move between MGEs by recombination.

## DISCUSSION

Here we describe VICSIN, a tool for the prediction and networking of integrated MGEs. VICSIN uses a consensus approach to MGE prediction that results in greater sensitivity than its constituent tools. It is especially powerful for poorly-studied host genomes such as those of the family *Bacteroidaceae*. Our data suggests that this may be a result of these tools being developed and tested using extant model organisms like those of *P. aeruginosa* and *E. coli* (18). VICSIN then uses nucleotide similarity to generate networks and clusters of the MGE predictions as a basis for classifying the MGEs and visualizing their relatedness. These networks provide insight into the frequency and routes of HGT and recombination occurring among MGEs. VICSIN analyses provide an import framework for understanding the variable fraction of microbial genomes.

We applied VICSIN to 88 *Bacteroidaceae* genomes from two groups within the family: (i) *B. fragilis* and (ii) *P. vulgatus* and *P. dorei. Bacteroidaceae* genomes are highly plastic, with large portions of these genomes predicted to be mobilized by HGT (66, 71–73). Although the mechanisms causing the entirety of these events are largely unknown, one of the principle drivers of HGT is likely uncharacterized integrated MGEs. Presently, between the NCBI and ICEberg databases only have 27 annotated *Bacteroidaceae*-specific phage or ICE elements. Our results found that 60% (57/95) of VICSIN-predicted MGE clusters lack a known MGE. Further, we show evidence of recombination within and between clusters of predicted MGEs, and hypothesize at least some of these MGEs are capable of transfer across a broad host range. It has been hypothesized that a combination of these factors is an especially potent driver of HGT across the *Bacteroidaceae* family (71, 73).

We found very limited gene sharing between known phages and integrated MGE predictions in the current *Bacteroidaceae* dataset. This suggests some barrier to recombination exists between some MGEs. We do not believe the barrier is caused by short infection times for these phages; Hankyphages are Mu-like viruses capable of stable integration into host chromosomes (74), and cells infected with CrAssphages have been observed to enter a “carrier state” where lysis and progeny release can be delayed for long periods of time (75). Part of this could be caused by structural constraints of phages versus ICEs. Whereas phages have limited space within the capsid to store their genome, ICEs which transfer directly from cell to cell via the T4SS have fewer constraints on genome size. Known ICEs can be hundreds of kilobases in length (*e*.*g*. PAIst in *Streptomyces turgidiscabies* >674 Kb) (76). Further, the current data reinforces the view that ICEs are likely the primary drivers of HGT in these genomes, though this may be due to a lack of knowledge about lysogenic phages infecting *Bacteroidaceae* hosts. Phages are known to drive HGT in other bacterial systems (77), so it stands to reason they should also play a role in *Bacteroides* and *Phocaeicola*.

Despite the limited gene sharing between phages and other MGE clusters, gene sharing between phages of the same cluster was found to be very high, covering the majority of the genomes examined. This further suggests phage genomes are under strong selection to only encode the content necessary for their transfer, to the exclusion of a diversity of accessory genes, including ARGs and T6SSs.

## METHODS

### Data availability

VICSIN, genome_grouper.py, and test datasets are available at github.com/IGBIllinois/VICSIN.

### Benchmark datasets

Two datasets were created and used for testing VICSIN, one of *Pseudomonas aeruginosa* genomes and another of *Bacteroides fragilis* genomes. Genomes in benchmark datasets were annotated based on published MGEs integrated in those genomes, or through manual annotation (Table S1).

The *P. aeruginosa* dataset consists of six ingroup genomes: *P. aeruginosa* DK2 (NC_018080), *P. aeruginosa* UCBPP-PA14 (NC_008463), *P. aeruginosa* PAO1 (NC_002516), *P. aeruginosa* LESB58 (NC_011770), *P. aeruginosa* PA38182 (HG530068), *P. aeruginosa* RP73 (NC_021577); and three outgroup genomes: *P. denitrificans* ATCC 13867 (NC_020829), *P. stutzeri* CGMCC 1.1803 (NC_015740), and *P. putida* KT2440 (NC_002947).

The *B. fragilis* dataset consists of six ingroup genomes: *B. fragilis* CL03T12C07 (NZ_AGXL00000000), *B. fragilis* NCTC 9343 (NC_003228), *B*. sp. 3_2_5 (NZ_ACIB00000000), *B. fragilis* 638R (NC_016776), *B. fragilis* CL05T00C42 (NZ_AGXO00000000), *B. fragilis* YCH46 (NC_006347); and three outgroup genomes: *B. fragilis* 3_1_12 (NZ_ABZX00000000), *B. nordii* CL02T12C05 (NZ_AGXS00000000), *B. salyersiae* CL02T12C01 (NZ_AGXV00000000).

### Viral, Integrative, & Conjugative Sequence Identification & Networking (VICSIN)

VICSIN takes genomes in nucleotide FASTA format or GenBank format. General Feature Format (GFF) files defining ORFs are optional inputs. In the absence of GFF or GenBank files, VICSIN uses ORFs called by VirSorter or Prodigal in later steps. Multiple genomes are input simultaneously and should represent a group of closely-related organisms. VICSIN performs an initial alignment of all genomes with Spine using default parameters to define a core genome and calculate Spine-PLA. If the Spine-PLA is less than 70%, a warning is passed to the user.

Prediction of MGEs by VICSIN is accomplished by three default tools (PhiSpy, VirSorter, AGEnt) and two optional BLAST-based searches (Known BLAST and CRISPR BLAST). VICSIN runs PhiSpy on each individual genome in a genome group with a window size (w) of 40 and threshold (n) of 20. VICSIN implements VirSorter with default parameters and the provided “Viromes” search database. Spine and AGEnt are run in sequence using default parameters except using a minimum core length of 500 bp. Known BLAST is done by default nucleotide BLAST using a user-input FASTA file of relevant known MGEs. CRISPR BLAST is based on CLdb (78). VICSIN finds overlapping predictions and merges them when they are within 5 bp of each other or a contig boundary. VICSIN breaks PhiSpy predictions at overlapping prediction boundaries to limit overprediction by PhiSpy. Finally, VICSIN uses Re-BLAST to compare all predictions, and will extend prediction boundaries to match those of other predictions made using multiple methods.

After prediction, VICSIN filters out predictions shorter than 12 kb, a cutoff chosen because this was the length of the shortest known MGE in the benchmark datasets; neither VICSIN or its constituent tools are optimized for the prediction of short MGEs, like insertion sequences. Next, VICSIN performs reciprocal nucleotide BLAST to compare all predictions to each other, and calculates total PLA for every prediction pair, which serves as the basis for networking. PLAs greater than or equal to 20% are passed to MCL with an inflation parameter of 2, generating MGE clusters. Predictions smaller than 12 kb are clustered separately.

### Other networking methods

vConTACT was run on CyVerse using default parameters except where noted. ClustAGE was run with default parameters and visualized with http://vfsmspineagent.fsm.northwestern.edu/results/cpQeC6nUglST/results.html. All networks were visualized with Cytoscape.

### Bacteroidaceae *experimental dataset*

From the NCBI web interface, 300 *Bacteroidaceae* genomes were downloaded (Table S2). An additional 18 genomes were added to the dataset that were sequenced as part of this work (Table S7). Known MGEs integrated in the networking steps were downloaded from NCBI or ICEberg databases (31).

### MGE cluster typing

Open reading frames from MGE prediction were annotated with the Pfam 33.1 database (79) and hmmscan using trusted cutoffs (--cut_tc). A list of Pfam protein families associated with ICEs and general mobile sequences was compiled (Table S5). Phage-associated Pfams were defined as any with the words “terminase,” “portal,” “capsid,” “tail,” “base plate,” “spike,” “neck,” “head,” or “phage protein” in their description. The number of phage, ICE, and mobile genes were counted. MGEs with more phage genes than ICE genes were assigned to the phage type. MGE MGEs with more ICE genes than phage genes were assigned to the ICE type. MGEs with no ICE or phage genes, or an equal number of phage and ICE genes, were assigned to the “mobile” type. Any MGE with no genes of interest were assigned to the “unknown” type.

### Gene annotation

Using the Pfam predictions from above additional Pfam IDs were assigned to functional categories based on gene annotations from *B. thetaiotaomicron* VPI-5482 and *B. fragilis* NCTC 9343 (Table S5). This included genes involved in capsule biosynthesis (CPS), polysaccharide utilization loci (PULs) and type VI secretion systems (T6SS). In addition, antibiotic resistance gene nucleotide sequences from the Comprehensive Antibiotic Resistance Database v1 (*n* = 2,163; https://card.mcmaster.ca/) (80) and an additional 17 genes missing from the database (*e*.*g*., *tetA, ermA, tetC*) were grouped into 90% identity clusters with USEARCH v5.2.236 (81). A BLAST database was then generated with the protein sequences for the 734 representative (seed) antibiotic resistance genes for these clusters. ORFs from the VICSIN predictions were then queried using an E-value cut off of 10^−10^, and the results were filtered for matches ≥ 50% aligned and ≥ 50% AA identity.

A BLAST database was generated with the protein sequences for 46 representative Hcp, TssB, and VgrG from *Bacteroidaceae* genomes (Table S6). ORFs from the VICSIN predictions were then queried using an E-value cut off of 10 ^-10^, and the results were filtered for matches ≥ 50% aligned and ≥ 50% AA identity.

### Genome sequencing

Cells were pelleted from 5 mL overnight culture in TYG (82) by centrifugation at 4,000 × *g* for 5 min at 4°C, resuspended in 0.5 mL TE buffer (10 mM Tris, 1 mM EDTA), and lysed by adding sodium dodecyl sulfate (SDS) and proteinase K (GoldBio, Olivette, MO) to final concentrations of 0.07% and 300 μg/mL, respectively, and incubating for 2 hr at 55°C. Cellular material was removed by washing twice in an equal volume of buffered phenol, phenol-chloroform-isoamyl alcohol (VWR, Radnor, PA), and DNA precipitated with 100% ethanol in the presence of 0.3M sodium acetate at -20°C overnight. DNA pellets were washed with 70% ethanol, dried, and resuspended in TE buffer.

DNA libraries were constructed with the Nextera XT Library Preparation Kit and Index Kit (Illumina, San Diego, CA). DNA libraries were pooled and sequenced on both the Illumina HiSeq5000 and HiSeq2500 and fastq files were generated from demultiplexed reads with bcl2fastq Conversion Software (Illumina, San Diego, CA). Reads were trimmed and assembled using the A5ud pipeline (83). Sequencing methods and assembly data are summarized in Table S7.

## ACKNOWLEDGEMENTS

We thank Ted Pacyga for his invaluable help with the genome_grouper.py program, Nadja Shoemaker and Abigail Salyers for access to an impressive collection of *Bacteroidaceae* isolates, and Alvaro Hernandez and Chris Wright for DNA sequencing.

This research received no specific grant from any funding agency in the public, commercial, or not-for-profit sectors. D.E.C. was supported by the University of Illinois at Urbana-Champaign Department of Microbiology and the NIH grant T32 DK077653-29. W.E.E. was supported by a James R. Beck Graduate Research Fellowship and NIH training grant T32 AI078876. R.J.W. was supported by the Allen Foundation with an Allen Distinguished Investigator Award.

